# Space-time dynamics of genome replication studied with super-resolved microscopy

**DOI:** 10.1101/2024.03.22.586000

**Authors:** Márton Gelléri, Michael Sterr, Hilmar Strickfaden, Christoph Cremer, Thomas Cremer, Marion Cremer

**Author notes:** corresponding authors Marton Gelleri; Thomas Cremer; Marion Cremer.

## Abstract

Genome replication requires duplication of the complete set of DNA sequences together with nucleosomes and epigenetic signatures. Notwithstanding profound knowledge on mechanistic details of DNA replication, major problems of genome replication have remained unresolved. In this perspective article, we consider the accessibility of replication machines to all DNA sequences in due course, the maintenance of functionally important positional and structural features of chromatid domains during replication, and the rapid transition of CTs into prophase chromosomes with two chromatids. We illustrate this problem with EdU pulse-labeling (10 min) and chase experiments (80 min) performed with mouse myeloblast cells. Following light optical serial sectioning of nuclei with 3D structured illumination microscopy (SIM), seven DNA intensity classes were distinguished as proxies for increasing DNA compaction. In nuclei of cells fixed immediately after the pulse-label, we observed a relative under-representation of EdU-labeled DNA in low DNA density classes, representing the active nuclear compartment (ANC), and an over-representation in high density classes representing the inactive nuclear compartment (INC). Cells fixed after the chase revealed an even more pronounced shift to high DNA intensity classes. This finding contrasts with previous studies of the transcriptional topography demonstrating an under-representation of epigenetic signatures for active chromatin and RNAPII in high DNA intensity classes and their over-representation in low density classes. We discuss these findings in the light of current models viewing CDs either as structural chromatin frameworks or as phase-separated droplets, as well as methodological limitations that currently prevent an integration of this contrasting evidence for the spatial nuclear topography of replication and transcription into a common framework of the dynamic nuclear architecture.

## Introduction

### The era of the 4D nucleome research

When the first, still unfinished reports of human genome sequencing were published in 2001, little was known about how the space-time dynamics of nuclear organization influences the gene expression patterns of different cell types ^1^. During the last two decades, Hi-C together with a wealth of other molecular biological tools as well as advanced microscopy, have led into the era of 4D nucleome research ^2–8^. This research has revealed an intricate space-time organization of nuclei with chromosome territories (CTs) built up from chromatin domains (CDs) with a DNA content in the order of 1 Mb. However, we still do not understand, how the 4D nucleome dynamically interacts from the level of the smallest chromatin structures to the level of entire CTs and possibly even larger networks of interacting CTs and how it is maintained during genome replication in cycling cells. In addition to epigenetic modifications, spatiotemporal changes of chromatin compaction and arrangements play a crucial role in cell type-specific gene expression patterns and other nuclear functions.

### DNA replication, a key process of nuclear function

In cycling cells, this functional 4D architecture must be transmitted from one cell cycle to the next. Replication of CTs with a single chain of CDs present in G1 yields CTs with two sister chromatids in G2. The resulting spacetime architecture of G2-CTs must allow the seemingly instantaneous formation of prophase chromosomes with two chromatids and the formation of daughter nuclei, preserving all functionally important structural features of the mother nucleus notwithstanding modifications necessitated by cell differentiation. Many details and mechanisms of how this is accomplished, have remained elusive.

At the molecular level, extensive knowledge has been gained on the replication process ^9–13^. Already in G1, accumulating prereplication complexes mark about 30.000 – 50.000 replication origin sites in mammalian cell nuclei, allowing the gradual accumulation of interacting components at replication origin sites ^14^. This gradual assembly yields functional replisome complexes during S-phase. Replisomes are complex macromolecular machineries ^15^ where an individual replisome controls and accomplishes DNA replication in discrete units (replicons) with a mean length of 175kb (Fig. 1A), for review see ^9,10,12,13^. Microscopic studies have essentially contributed to the current state of knowledge on replication ^16^. For instance, early studies on stretched DNA fibers, prepared from nuclei after pulse-replication labeling, revealed that several adjacent replicons are synchronously activated resulting in replication domains (RDs) with a length of ~0.5 – 1Mb ^17^ (Fig. 1B). Several RDs can be arranged in blocks up to ~10 Mb with similar replication times ^18^.

**Fig. 1.**
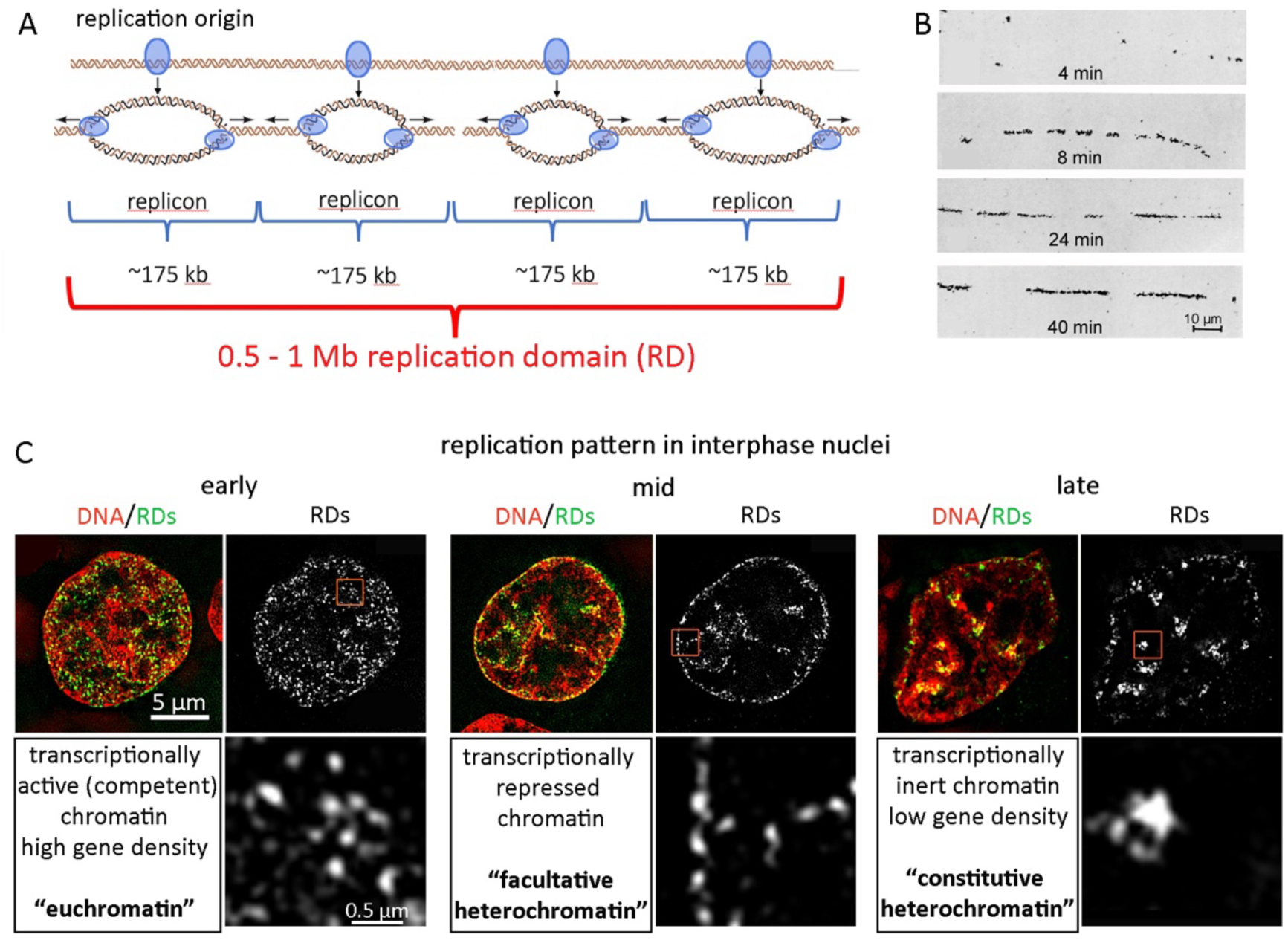
DNA replication in one and three dimensions. **(A)** Simplified scheme of DNA replication along the DNA strand starting with the activation of the replication machinery (replisome) at a replication origin site (marked as blue ellipsoids) with bidirectional unwinding and subsequent reduplication of the DNA strand. An individual replisome controls and accomplishes DNA replication in discrete units (replicons) with a mean length of ~175kb (for references see text). **(B)** Early microscopic evidence (“1D-microscopy”) for distinct replication sites and estimation of replication speed) on stretched DNA fibers after pulse labeling of CHO nuclei with ^3^H thymidine for different time length. The predicted calculated length of a stretched DNA fiber is 0.34 µm/kbp. Values for the 8, 24 and 40 min label times fit quite well with a replication speed of ~2kb/min (Fig. modified from ^17^ with permission). **(C)** Single optical sections of super-resolved images after EdU labeling obtained by 3D-structured illumination microscopy (3D-SIM). Typical replication patterns for early, mid and late replication in interphase nuclei demonstrates the tight coupling of replication time with genome architecture.

2D and 3D microscopic studies on replication labeled interphase nuclei demonstrated the genome-wide partitioning into discrete structural entities, called replication domains (RDs) with typical replication patterns for early, mid and late replication ^19^. These patterns have been evolutionary conserved ^20,21^ and indicate the tight coupling of replication timing with nuclear architecture (Fig. 1C). Globally, transcriptionally competent/active gene-rich domains in the nuclear interior replicate early, followed by (mid)-replication of more condensed chromatin domains harboring repressed DNA, preferentially located at the nuclear and nucleolar periphery. Heterochromatin blocks, largely constituted by repetitive sequences in clusters of pericentomeric heterochromatin (chromocenters), represent the major part of late replicating chromatin, for review see ^11,22^. RDs were microscopically defined as ~1Mb chromatin domains (CDs) ^23,24^ (see below for further details). They persist as stable chromatin units throughout interphase and during subsequent cell cycles. Using super-resolution microscopy, RDs can be optically resolved down to clusters of a few single replicons (150–200 kb) ^25,26^ or even smaller structures ^27,28^.

### Challenging issues of DNA replication in the context of a functional nuclear architecture

Fig 2A presents an overview on a recent study on true-to-scale DNA-density maps of chromatin where absolute chromatin compaction differences in the range between <5 Mb/μm3 up to >300 Mb/μm3 were described using single molecule localization microscopy (SMLM) ^29^. Such massive differences, if confirmed by future studies, have a strong impact on the accessibility of individual macromolecular complexes to their DNA target sites such as the replisome assembly ^30^. According to Maeshima and colleagues ^31,32^, nucleosome arrangements at a density <40 Mb/µm^3^ provide sufficient space to allow a high degree of accessibility for macromolecular protein complexes that have been typically assigned with a size >20nm diameter ^32,33^. Above this density (marked in brown color in Fig. 2A) their accessibility becomes severely constrained for individual macromolecules, and access of macromolecular aggregates formed outside is excluded, for review see ^34^.

**Fig. 2.**
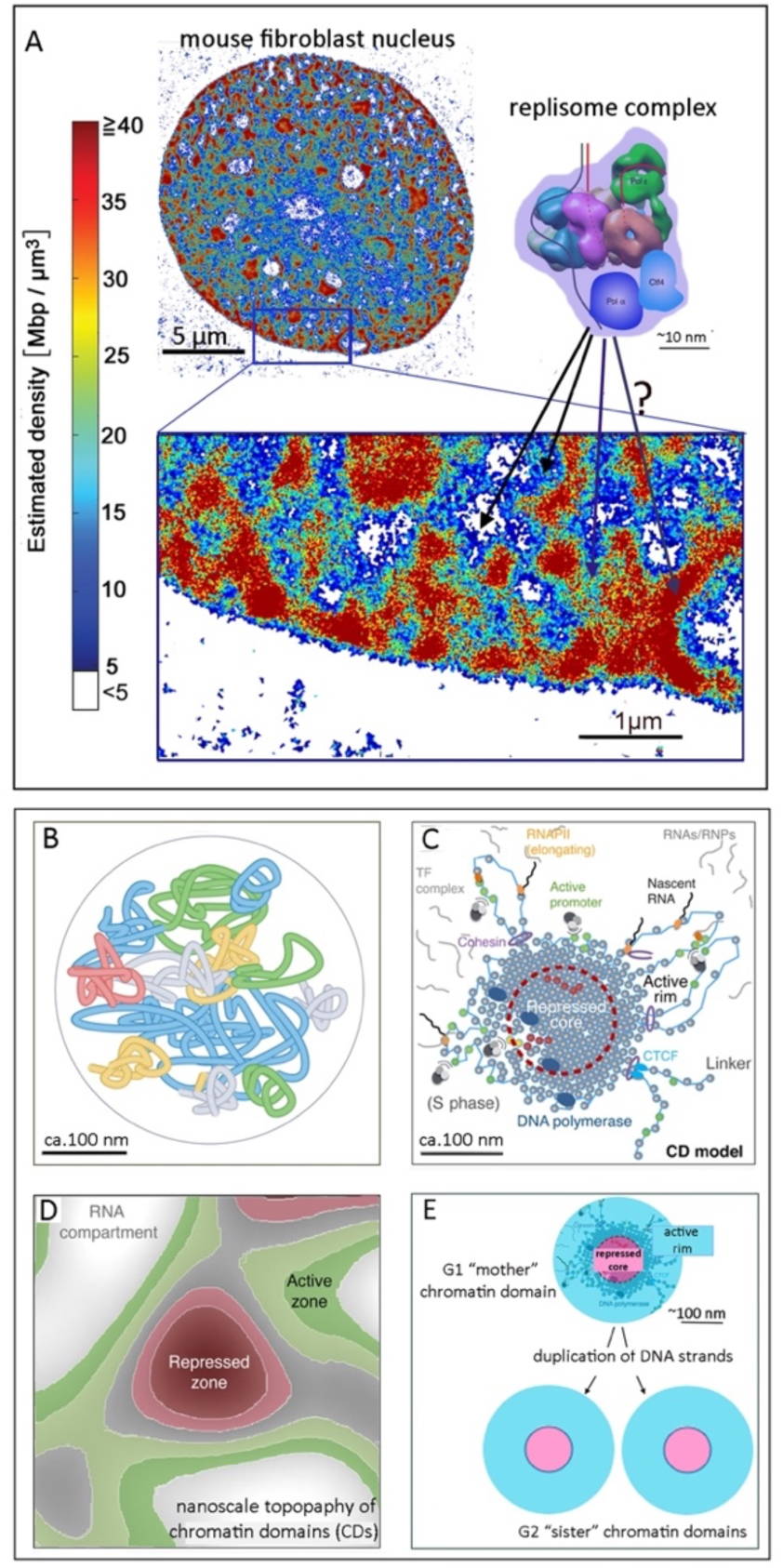
Problems of genome replication in the context of higher order chromatin landscapes. **(A)** 100 nm optical mid-section with 20 nm lateral resolution through a mouse fibroblast nucleus recorded with single molecule localization microscopy (SMLM), adopted from ^29^. DNA was stained with Sytox and color coded (see color bar code) for representation of absolute DNA densities noted within the ANC <40 Mb/µm^3^. Inset magnification of boxed area: white=interchromatin compartment (IC); blue/green/orange = perichromatin region (PR), assigned to the active nuclear compartment (ANC); dark red/brown = compact chromatin with densities >40 Mb/µm^3^ up to >300 Mb/µm^3^ (INC). **(B)** Cartoon of a topologically associated domain (TAD based on Hi-C) shows the folding of a contiguous chromatin fiber (modified with permission from Fig. 4 in ^69^). The access of individual components and large macromolecular complexes into the interior appears virtually unrestricted but may be constrained depending on the space-time dynamics of the fiber convolute (**C)** Cartoon of an ~1 Mb chromatin domain (CD) based on 3D-SIM data shows a repressed core with densely compacted nucleosomes indicated as little grey dots (INC) and a decondensed periphery (ANC) with transcriptionally active loops expanding from the compact core (adapted from Fig. 4 in ^37^ with permission). **(D)** Cartoon of the nanoscale topography of CDs embedded in the chromatin landscape (adapted from Fig.4 in^37^ with permission) indicates a zonal map of a mountainous chromatin landscape with a repressed zone in the center (INC). IC-channels and lacunae are lined by the PR. Together they constitute the transcriptionally active ANC. These pronounced differences of DNA compaction suggest constrained movements of replisome complexes in the ANC and exclusion from highly compacted chromatin segments **(E)** The ANC-INC model predicts that genome replication, in addition to the replication of DNA nucleosome and epigenetic markers, requires the faithful maintenance of the nuclear landscape. Accordingly, replication of a mother CD in G1 should yield two sister CDs during or after S-phase with the same inside-outside structure. The underlying mechanisms have remained elusive. In contrast, this problem would not exist for CD models with a random chromatin fiber arrangement or a droplet-like organization based on chromatin phase separation.

Fig. 2B-E show state-of-the-art models of chromatin landscapes and problems, which arise when we consider genome replication in the context of these models. Spatial access of replication machines may be taken for granted on the assumption of expanded chromatin loops (Fig. 2B), whereas the access to highly compacted chromatin provides a challenging problem (Fig. 2C).

Both microscopy and Hi-C studies demonstrated chromosome territories built up from ~1 Mb sized CDs or topologically associating domains (TADs) composed of multiple nucleosome clutches and chromatin nanodomains (CNDs) ^27,28^ as basic substructures of chromosome territories (CTs). The DNA content of microscopically defined CDs is comparable to estimates for TADs detected in Hi-C experiments. TADs and CDs/RDs are used synonymously ^13,35^. This equivalence, however, neglects the profound differences of both methods to describe different aspects of higher order chromatin structures above the level of individual nucleosome (for a critical comparison see also Supplementary Table 1 in^36^), namely folding of chromatin fibers, the superior task area of contact-matrix based Hi-C studies, and the quantitative analysis of DNA/chromatin compaction and intranuclear positions, the field of high-resolution microscopy.

Despite intense ongoing research, the space-time organization of CDs has remained a challenging problem. Evidence has been reported for both microphase separation of chromatin droplets ^31,35^ and for a defined structural organization^2,37^. Transitions between states of phase-separation and a defined structural organization seem possible ^38^. In this respect, the size of the investigated structures plays a decisive role. Whereas small CDs or smaller structures, such as chromatin nanodomains and nucleosome clutches can behave like chromatin droplets, a structural organization is evident for CTs and in particular for mitotic chromosomes, which require physical stiffness and rigidity for proper separation ^39^. Ronald Hancock pioneered research into the role of macromolecular crowding in the assembly and function of nuclear compartments ^40–42^. Arguably, in the presence of cations and a macromolecular crowder CDs can take on the state of liquid droplets that become stiffer, more solid-like and further condensed by additional crowders ^43^.

Our currently favored model argues for a well-defined structural organization of CDs with a compact core region present in the CD interior, attributable to the inactive nuclear compartment (INC) and a transcriptionally active periphery, the active nuclear compartment (ANC) ^2,37,44^ (Fig. 2C-D). Nuclear landscapes show a nanoscale zonal organization of CDs (Fig. 2D) with density peaks forming a highly compacted repressed core in the center, surrounded by a decondensed periphery, the perichromatin region (PR) ^45^ where active chromatin loops ^31,37^ expand into the lining interchromatin compartment (IC), a system of wider lacunae largely void of chromatin but containing nuclear speckles and bodies and of channels connected to nuclear pores ^36,46–48^. Microscopic studies support diameters of CDs between 200-400 nm ^37^. However, it is difficult to delineate individual CDs in the mountainous chromatin density landscape illustrated in (Fig. 2A). Caution is recommended when comparing such data from different studies, as measurements are strongly influenced by the inclusion of a decondensed periphery or of only the compact core region and can be influenced by fixation protocols. Furthermore, it is important to realize that liquid droplet models of CD organization and structural models have strongly different biophysical implications for CD duplication ^49^. For example, in a structured CD individual DNA sequences may be permanently located within the center of the compact core region. A positional change toward the periphery in order to form a transcriptionally competent loop may require a particular molecular mechanism. In a liquid droplet, by contrast, positions are permanently changeable. Accordingly, a sequence recorded at the center of the core region in a high-resolution live cell snap shot may be found soon after at its surface.

Fig. 2E illustrates implications of CD duplication based on our currently favored model. When a mother chromatid domain (mCD) present in CTs at G1 of the cell cycle yields two homologous sister chromatid domains (sCDs) during S-phase, functionally important structural features of the mCD must be maintained during replication or restored thereafter. How long this spatial reconstruction takes is not known. Problems of a special reconstruction are mitigated in droplet models of CDs. Here, the internal organization of CDs is based on inherent features of phase separation and does not require special mechanisms for movements of nucleosome clutches towards the CD periphery prior to replication and back into interior movement thereafter. Moreover, in G2-chromosome territories sCDs are arranged along the two newly formed sister chromatids. The actual spatial arrangements of these interphase sister chromatids are still not well understood, but they must meet the functional requirements to separate and form the two chromatids of prophase chromosomes at the onset of mitosis, as well as the formation of daughter nuclei, preserving functionally important structural features.

### Topography of DNA replication compared with transcription in the context of the ANC-INC model

Previously, our research team demonstrated that transcription occurs in the ANC ^36,44,47,48^. The experiments were performed with 3D structured illumination microscopy (3D-SIM) and quantitative image analysis of SIM serial sections of nuclei following immune-cytochemical detection of RNA Polymerase II (RNAPII) and epigenetic markers associated with transcriptionally competent or silent chromatin, respectively ^34^. Essential features of this approach and pertinent results are summarized in Fig. 3.

**Fig. 3.**
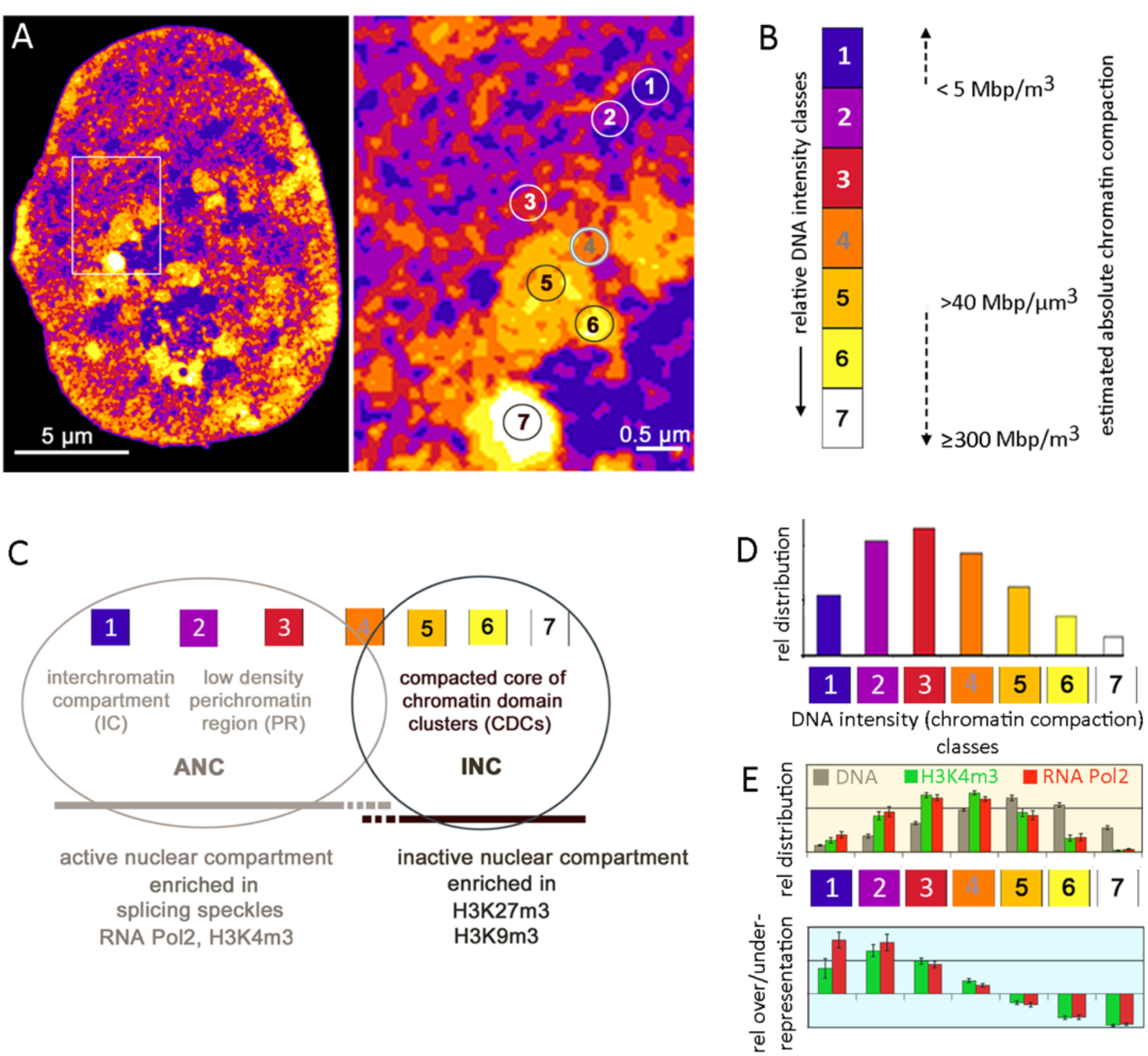
ANC-INC model and the topography of transcription. **(A)** Voxel-based classification of DAPI stained DNA into seven intensity classes with equal intensity variance as proxy for relative chromatin compaction differences visualized as color heat map in an optical SIM serial section of a mouse myoblast nucleus. Inset magnification with encircled areas in each class exemplifies chromatin domain clusters (CDCs) with a zonal organization of decondensed CDCs at the periphery adjacent to the interchromatin compartment (IC) and higher compacted chromatin located in the CDC interior. (**B**) Voxels attributed to color-coded intensity classes 1-7 (note the different color coding compared to the color coding shown in Fig. 2A for SMLM images). Voxels of classes 5-7with estimated intensities >40 Mb/µm^3^ (compare Fig. 2A) are attributed to the inactive nuclear compartment (INC), and voxels with intensities <40 Mb/µm^3^ to the active nuclear compartment (ANC), compare Fig.2A. **(C)** Schematic overview of functional assignment of DNA intensity (chromatin compaction) classes. The lowest DNA density class 1 represents the interchromatin compartment (IC), proposed as the preferred transport system for imported proteins, exported RNPs, and the formation of macromolecular aggregates. Intensity classes 2 – 3 comprise low-compacted chromatin lining the IC (perichromatin region, PR). The PR represents the periphery of chromatin domains (CDs) and chromatin domain clusters (CDCs) and is the preferential nuclear subcompartment for transcription. Class 4 denotes a transition zone, classes 5–7 comprise the compact chromatin of the INC. **(D)** Relative distribution of DNA density classes based on a quantitative 3D SIM analysis of the DAPI stained nucleus shown in A. **(E)** Example for the quantitative mapping of ‘active transcription’ marks H3K4m3 and RNAPII of a human hematopoietic progenitor cell on DNA intensity classes delineated as relative distribution (top) or as over/under-representation (bottom) in the respective intensity classes. The profiles show a clear enrichment of these marks in the low intensity classes 1-4 and an under-representation in classes 5-7 indicating that transcription occurs predominantly in the ANC (adapted from Fig. 5 in ^47^, for experimental details see ^50^).

In the present study, we used this approach to study the nuclear topography of replication domains (RDs). Cells were pulse-labeled for 10 min with EdU, and fixed either immediately or after a chase of 80 min, considered sufficient for the completion of DNA duplication of a pulse-labeled RD/CD. As our working hypothesis, we predicted to observe the same topography for DNA replication as previously found for transcription (Fig. 3E). Our results clearly argue against this hypothesis. We discuss the consequences of this rejection both with regard to methodological shortcomings of our preliminary study, in particular the resolution limit of 3D SIM and potential limits of the pulse-labeling (10 min) and chase period (~80 min), but also with regard to limitations of and contradictions between current models of CD/TAD organization.

## Materials & Methods

### Cells

PMI28 cells, a near-diploid mouse myoblast cell line, were grown in HAM-F10 Medium supplemented with 20% FCS and 1× Penicillin/Streptomycin. To prevent differentiation into myotubes, cells were constantly kept in subconfluent culture conditions. Mouse embryonic stem cells (mESCs), derived from the clone 16.7 of a *Mus musculus × Mus castaneus* hybrid female mESC line were grown in DMEM with GlutaMAX medium, supplemented with 16% FBS, 1× Non-essential amino acids, 0.1mM β-Mercaptoethanol. Maintenance of the undifferentiated state was achieved by adding 1000 U/ml LIF, 1μM PD 0325901, 1 μM CHIR 99021 to the medium. In addition, care was taken to thoroughly separate cells during the passage procedure and to prevent the formation of large colonies.

### Pulse replication-labeling of replication domains (RDs), fixation and DNA staining

For pulse-replication labeling cells were incubated in medium containing 5-Ethynyl-dU (EdU) at a final concentration of 10 μM for 10 min. After 10min incubation in EdU, cells were washed in 1xPBS, and fixed with 2% formaldehyde/PBS for 15min. Cells were washed in 1xPBS and incubated with 0.5 U/ml RNase A and 20 U/ml RNase T1 (Ambion, USA) for 1 h at 37°C, followed by permeabilization with 0.5% Triton X-100/PBS/ Tween 0.02% for 10 min. EdU detection was performed by a Cu(I) catalyzed cycloaddition reaction that covalently attaches a fluorescent dye (azide modified Atto655 for mouse myoblast cells, azide modified Alexa488 for mouse embryonic stem cells) to the ethynyl-group of the nucleotide according to manufactures instructions (baseclick). For cells with an additional chase after replication labeling, EdU containing medium was replaced after careful washing in 1xPBS by normal growth medium for 80 min before fixation. Mouse myoblasts were counterstained in Sytox Orange (0.5 mM), mESCs were stained in DAPI (5 mg/ml), postfixed in 4% formaldehyde/PBS for 10 min and mounted in antifade mounting medium (Vectashield).

### 3D-structured illumination microscopy (3D-SIM)

Super-resolution structured illumination imaging was performed on a Zeiss Elyra 7 microscope equipped with a 63x/1.46 alpha Plan-Apochromat oil immersion objective (Carl Zeiss, Jena). Raw data image stacks were acquired with 13 raw images per plane and an axial distance of 116 nm. Dual color images were recorded sequentially. 561 nm and 640 nm lasers were guided by a quadband reflector module (LBF 405/488/561/642) to the sample. Fluorescence was collected and then split by a Duolink filter (SBS BP490 – 560 + LP650, Carl Zeiss, Jena) onto two cameras. Atto655 was excited using the 640 nm laser and wavelengths above 650 nm were imaged on camera 1, and Sytox Orange fluorescence was excited by the 561 nm Laser and wavelengths between 560 nm and 640 nm were collected on camera 2. SIM image reconstruction was done in the Zeiss Zen Black software.

For replication studies of chromocenters shown in Fig. 6, 3D-SIM was carried out using an OMX Version 3 (Applied Precision) microscope equipped with a UPlan APO 100× 1.4 NA oil-immersion objective (Olympus), illumination lasers for 405 and 488 nm and Photometrics Cascade II EMCCD cameras for the respective color channels. Samples were screened on a personal Delta Vision (pDV, Applied Precision) microscope, a conventional wide field microscope equipped with a motorized stage, a 60× 1.4 NA oil-immersion objective, a Xe-illumination and a Photometrics Cool-Snap camera. For each image taken with the pDV, the position of the motorized stage, which is compatible to the OMX stage, was saved. Samples were mounted on the OMX (1.512 oil), stage coordinates were transferred and the respective positions were imaged. According to the size of the nucleus depicted, image stacks with a z-distance of 0.125 μm were acquired at a resolution of 512 × 512 or 256 × 256 pixels, corresponding to an imaging area of 40 × 40 μm or 20 × 20 μm, respectively (acquisition software: DeltaVisionOMX Vers. 2.25). The image raw data was subsequently reconstructed to high resolution images using the SoftWorx Vers. 5.1.0 software. Final processing (i.e. thresholding and composition) of reconstructed images was done in Fiji (ImageJ) and Photoshop.

### 3D assessment of Sytox intensity classes as proxy for chromatin compaction classification

Nuclei voxels were identified automatically from the Sytox channel (560 nm – 640 nm) intensities using Gaussian filtering and a fixed threshold of 0.003. We found that due to different noise levels the fixed threshold was more reliable than an automatic threshold determination. For chromatin quantification a 3D mask was generated in R to define the nuclear space considered for the segmentation of Sytox signals into seven classes with equal intensity variance by a previously described in house algorithm ^50^ available on request. In brief, a hidden Markov random field model classification was used, combining a finite Gaussian mixture model with a spatial model (Potts model), implemented in the statistics software R ^51^. This approach allows threshold-independent signal intensity classification at the voxel level, based on the intensity of an individual voxel. Color or gray value heat maps of the seven intensity classes in individual nuclei were generated in ImageJ.

### Semi-automatic segmentation of EdU-labeled signals and their quantitative allocation on 3D chromatin compaction (Sytox intensity) classes

Individual voxels of EdU signals of the respective marker channel were segmented using a semi-automatic thresholding algorithm (using custom-built scripts for the open-source statistical software R http://www.r-project. org, available on request). Xyz-coordinates of segmented voxels were mapped to the seven DNA intensity classes. The relative frequency of intensity weighted signals mapped on each Sytox intensity class was used to calculate the relative distribution of signals over chromatin classes. For each studied nucleus, the total number of voxels counted for each intensity class and the total number of voxels identified for the respective EdU signals were set to 1. As an estimate of over/under representations (relative depletion/enrichment) of marker signals in the respective intensity classes, we calculated the difference between the fraction of voxels calculated for the EdU signals and the corresponding percentage points obtained for the fraction of voxels for a given Sytox intensity class. This difference was then normalized (oder divided by?) to the respective fraction of voxels in a given Sytox class. Calculations of Sytox intensity classes and relative distributions of EdU signals were performed on single-cell level. Average values over all nuclei were used for evaluation of the relative depletion and enrichment of EdU signals and plotting. For a detailed description, see ^47,50^.

### Calculation of ratio images

Ratio images were calculated using Fiji. First, the mask of the cell nucleus as determined under “*3D assessment of Sytox intensity classes as proxy for chromatin compaction classification*” was applied to both the SYTOX Orange SIM images and the EdU SIM images. The resulting z-stacks contained the total Sytox and total EdU signal intensities inside the 3D cell nucleus. The sum intensity across the 3D stacks yields then the total intensities of SYTOX Orange and EdU channels corresponding to the total amount of SYTOX bound to DNA and the total EdU signals within the cell nucleus. We then divided for each channel every image of the masked 3D SIM stack by the respective sum intensity. For the SYTOX image, each pixel intensity in such a normalized image then corresponds to the fraction of DNA bound SYTOX molecules, and for the EdU image, each pixel corresponds to the fraction of EdU signals. EdU to SYTOX ratio images were calculated by dividing the normalized EdU images by the normalized SYTOX images. The ratio R_i_ in a pixel i is given by

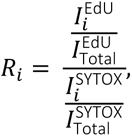

with I^EdU^_i_ being the intensity in pixel i of the EdU channel, I^SYTOX^_i_ being the intensity of pixel i in the SYTOX channel, and I^EdU^_Total_ and I^SYTOX^_Total_ being the total intensity in the EdU and SYTOX channel respectively.

## Results

Using 3D-SIM, we addressed the topography of DNA replication in the context of the ANC-INC model with the following questions: are active replication sites preferentially seen in decondensed chromatin classes assigned to the ANC, similar to transcriptionally active/competent chromatin? To what extent differs the distribution of labeled RDs during ongoing replication from RDs after accomplished replication indicating a dynamic process of chromatin during/after replication?

Nuclei of mouse myoblasts (Pmi28 cells) were pulse-replication labeled for 10 min with EdU and fixed either immediately thereafter or after an additional chase of 80 min respectively. These two time points were chosen for the following rationale: replication in mammalians proceeds with a speed of about 2-3kb/min along the DNA strand ^12^ where one replicon (the DNA segment replicated by a single initiation site) covers on average 150-200kb ^25^. As 3-6 adjacent replicons typically fire synchronously and form 0.5 - 1Mb replication domains ^11,52^ (see also Fig. 1), the full replication of an entire RD takes about 1h. In reverse, within a 10 min replication pulse about 10-20% of an individual RD are labeled. This approach yields >1000 3D-SIM detectable EdU foci at a given time point (data not shown, compare ^25,36,37^). Within a subsequent 80min chase the replication of an entire ~1Mb RD should be fully accomplished, accordingly this condition represents the post-replication status of a labeled RD. The replication of clustered replication foci in large heterochromatin blocks, constituting chromocenters of several Mb in mouse nuclei, may take longer ^53,54^.

Nuclei were counterstained with Sytox Orange, a fluorescent DNA dye without DNA sequence binding preference ^44^. Optical serial sections of labeled nuclei with either replicating CDs (10min EdU pulse) or replicated CDs (10min EdU pulse + 80min chase) state were categorized visually with either a typical early or late replication pattern. (Figs. 4 and 5). The respective inset magnifications of both series in nuclei with an early replication pattern showed replication foci covering all density classes on respective heatmaps, with labeled segments often expanding over several classes (Figs. 4A-B). In nuclei after an 80 min chase (Fig. 4B) the preferential localization of EdU labeled sites in the small compacted island-like cores which are part of early replicating ‘euchromatic’ regions is of note.

**Fig. 4.**
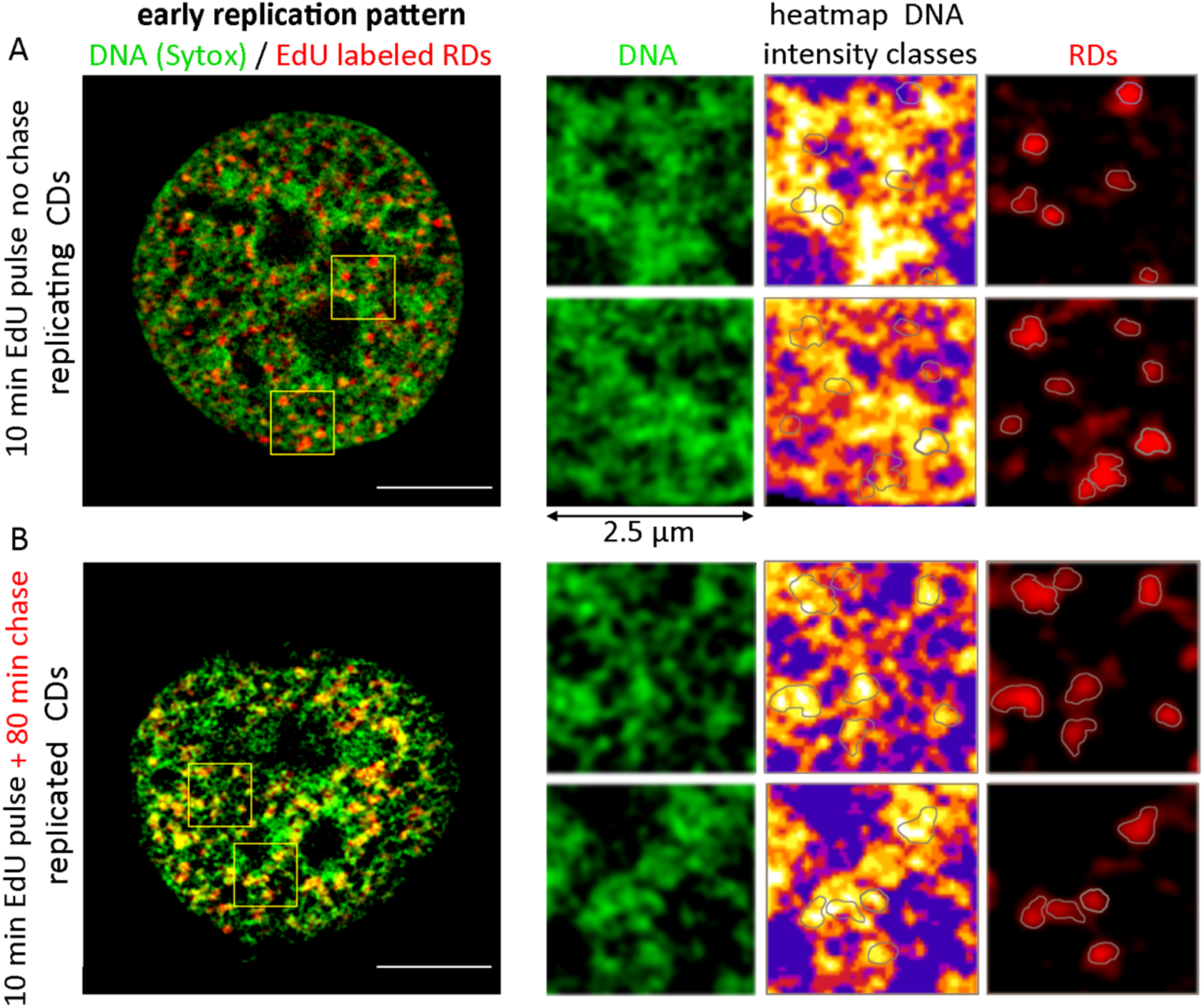
Light optical mid-sections from whole 3D-SIM acquisitions of nuclei with an early replication pattern. **(A)** Nucleus fixed after 10 min EdU pulse labeling with replicating RDs/CDs. **(B)** Nucleus fixed after an 80 min chase shows replicated CDs. Representative inset magnifications show an overlay of labeled RD borders on DNA intensity heatmaps (left: DNA staining, middle: corresponding heatmap of DNA intensity classes, right: labeled RDs). Labeled RDs are seen both at decondensed chromatin sites and in or at the surface of compacted chromatin domain clusters. Note the localization of replicated CDs in small condensed heterochromatic islets throughout early replicating euchromatic regions (size bars = 5 µm). 3D-SIM acquisitions of the whole nuclei are provided in Supplemental Movies S_movie_1and 2.

**Fig. 5.**
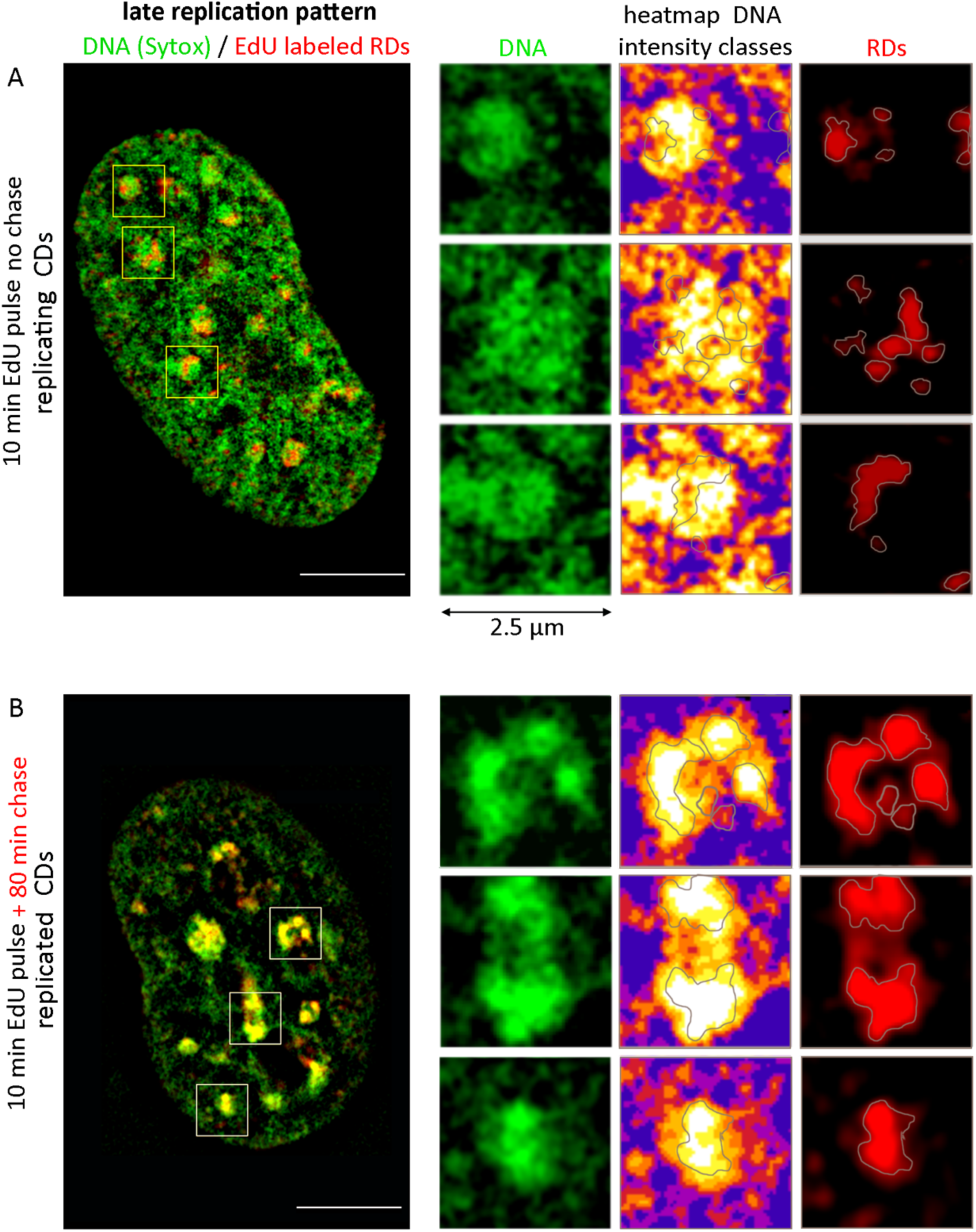
Light optical mid-sections from whole 3D-SIM acquisitions of nuclei with a late replication pattern. **(A)** Nucleus fixed after 10 min EdU pulse labeling with replicating RDs/CDs. **(B)** Nucleus with replicated CDs fixed after an 80 min chase. Representative inset magnifications show late replicating chromocenters (left: DNA staining, middle: corresponding heatmap of DNA intensity classes, right: labeled RDs/CDs). Replicating CDs (A) show the preferential localization of labeled RDs in decondensed chromatin, extending into or through chromocenters. After the chase (B), fully replicated CDs are seen within highly condensed chromatin clusters (size bars = 5 µm). 3D-SIM acquisitions of the whole nuclei are provided in Supplemental Movies S_movie_3-4.

The visual impression in nuclei with a late replication pattern, which is essentially, though not exclusively characterized by the replication of large heterochromatin blocks (chromocenters) differed remarkably between both series: as exemplified in Fig. 5, in cells with replicating CDs (10 min pulse) EdU signals were preferentially located in regions of decreased chromatin compaction permeating the chromocenters (Fig. 5A). In contrast, nuclei with replicated CDs recorded after the 80 min chase (Fig. 5B) revealed labeled DNA segments mostly in highly compacted clusters of chromocenters. 3D-stacks of whole nuclei are provided in supplementary movies S1-4.

For comparison, a series of EdU-replication patterns of chromocenters in undifferentiated female mouse embryonic stem cells (mESCs), pulse-labeled with EdU for 10 min, and fixed immediately after the pulse or after an additional chase of 20, 40, 60 and 80 min, is provided in Fig. 6. In these cells the width of major IC-channnels pervading the chromocenters appears wider compared to the somatic cells shown in Fig. 5. This experiment illustrates the location of replicating DNA located both within and at the edge of wide IC-channels pervading the interior of chromocenters in the first three time points (0, 20, 40 min chase), followed by a subsequent relocation into the lining heterochromatin (60, 80 min chase).

**Fig. 6.**
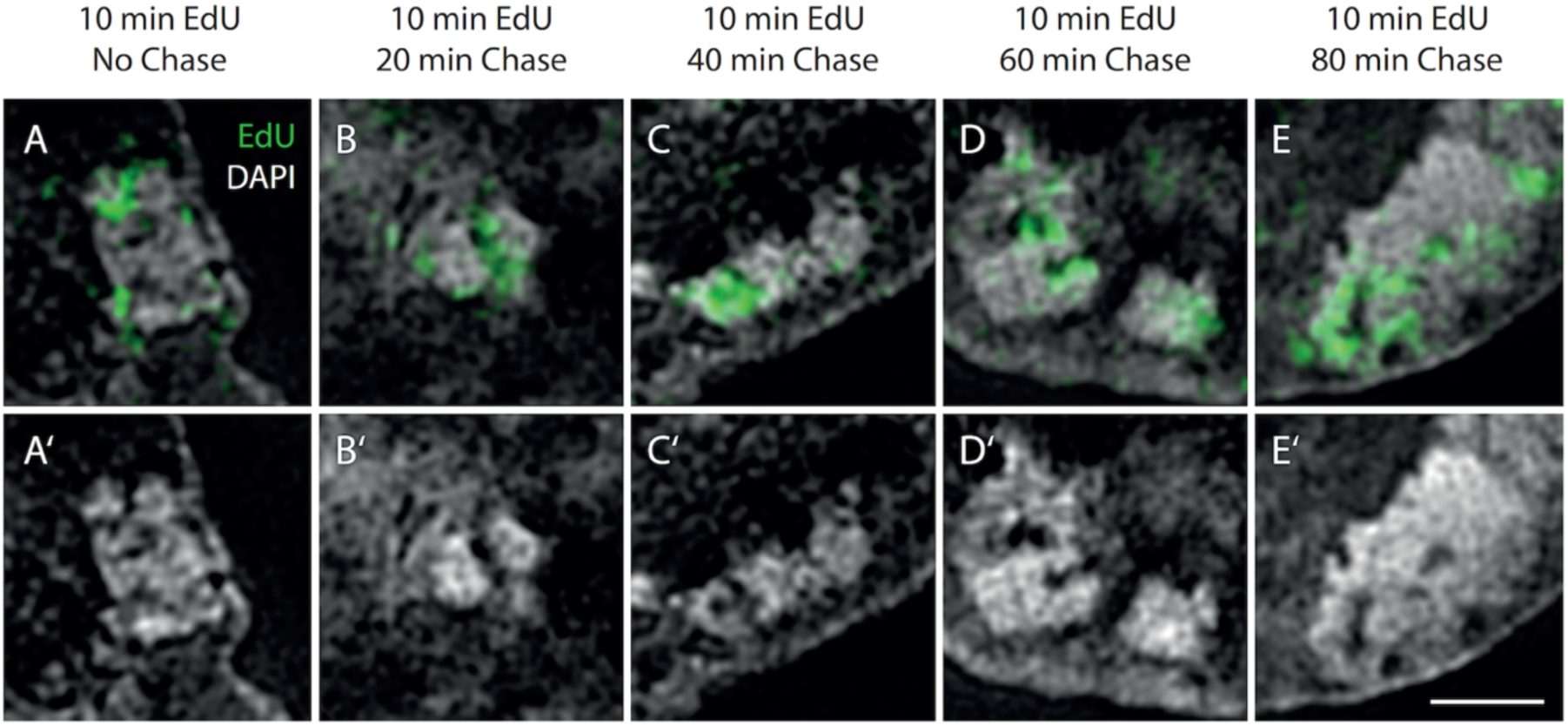
EdU-Replication Patterns of DAPI stained chromocenters in undifferentiated mouse female ESCs. Cells were pulse-labeled with EdU (green) for 10 min, and fixed immediately after the pulse (panel A) or after an additional chase of 20, 40, 60 and 80 min (panels B-E). For comparison, panels A’–E’ show the DAPI stained chromocenters alone (grey). (bar = 1 μm). This experiment illustrates the location of replicating DNA both within and at the edge of wide IC-channels pervading the interior of large heterochromatin blocks, called chromocenters after a 10 min EdU pulse and chase periods of 20 and 40 min (evident in panels A-C), followed by a clearly recognizable relocation into the lining heterochromatin during longer chase periods (D and E). Taken from unpublished images from Michael Sterr, diploma thesis, “Three Dimensional Structured Illumination Microscopy Studies of the Nuclear Architecture in Murine Embryonic Stem Cell and Preimplantation Embryos”, LMU 2012.

For the mouse myoblast (Pmi28) cells a 3D quantitative assessment of the distribution of EdU-labeled signals over the seven DNA intensity classes was performed from five nuclei of each series. The delineation as relative over- or underrepresentation in each DNA density class confirmed their representation in all classes (Fig. 7, compare also Fig. 3 E). For early replicating DNA (Fig. 7A, blue columns) these plots reveal a slight underrepresentation in classes 1-2 and an overrepresentation in classes 4-6 (marginal in class 7) in nuclei fixed after the 10 min pulse. In contrast, after the additional 80min chase the replicated DNA was found strongly overrepresented in classes 5-7 (grey columns in Fig. 7A), with the most prominent increase in class 7, the latter, however, represents only a small fraction (<5%) of the total DNA. This shift toward high intensity classes indicates a dynamic behavior of early replicating chromatin during the chase. Quantitative assessment of late replicating chromatin (Fig. 7B) shows essentially similar results, however, the overrepresentation of late replicating DNA in intensity classes 6-7 is higher (Fig. 7B, brown bars), in agreement that late replicating DNA mainly comprises pericentromeric heterochromatin. This over-representation becomes distinctly more pronounced after the 80 min chase. The distribution of EdU labeling clearly refutes our prediction that labeling should be predominantly observed in low DNA intensity classes.

**Fig. 7.**
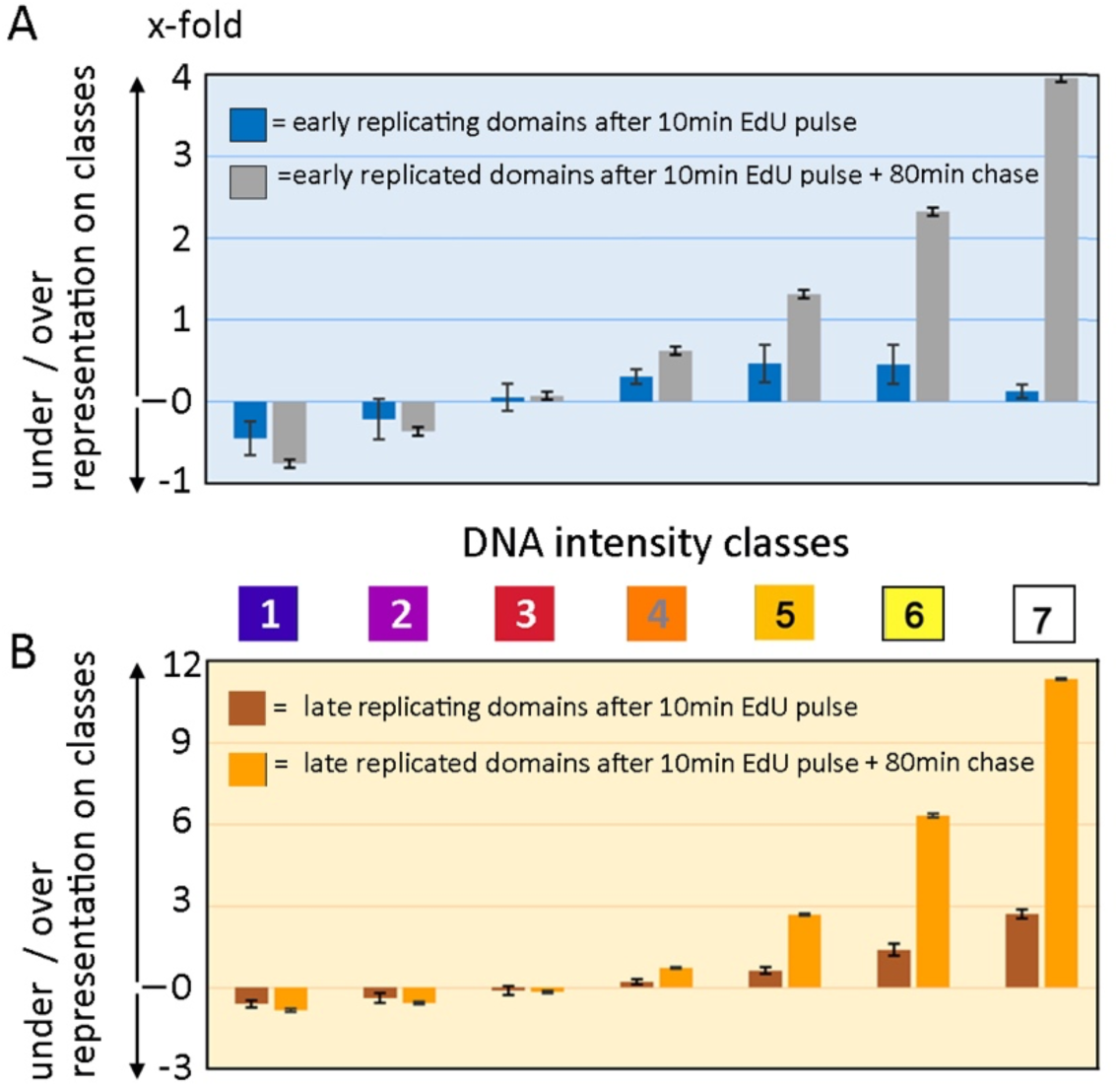
Relative representation of EdU-labeled RDs on respective DNA intensity classes in Pmi28 mouse myoblasts. Quantified levels of relative enrichment/over-representation (positive values) or depletion/under-representation (negative values) of labeled RDs in the seven chromatin compaction classes plotted as x-fold excess. **(A)** Early RDs after 10min pulse labeling (blue) and after 80min chase (grey). **(B)** Late RDs after 10min pulse labeling (brown) and after 80min chase (orange). An over-representation in classes 5-7 is higher in RDs recorded after an 80min chase, both for early and late replicating chromatin. For each series 5 nuclei (each comprising between 30-40 optical sections were evaluated. Note the different scales in Y-axis for “fold increase”. Error bars delineate standard deviations.

Evidence for a dynamic behavior of pulse-labeled chromatin during the chase period is supported by color-coded ratio images of normalized EdU signals and DNA signal intensities (Fig. 8). 3D-stacks of whole nuclei are provided in supplementary movies S_5-8. In case of a rapid redistribution of replicated chromatin we would expect ratios close to 1 (dark purple) indicating that the amount of replicated chromatin reflects the DNA density at a given site. Instead, we observed in nuclei with replicating domains (Fig. 8 left panels, in A and B) still big clusters with increased ratios >4 (yellow/white) indicating an excess of labeled chromatin surrounded by larger EdU-labeled regions with ratios of 1-3 (purple-red). In nuclei studied after the 80 min chase (Fig. 8 right panels in A-B) purple/red colored regions are more prominent at the cost of the yellow/white clusters suggesting a more uniform distribution of the EdU signal. Notably, distinct white dots mark preferably the periphery of EdU-labeled early and late replicated regions indicating that the process of the redistribution of replicated chromatin was not completed during the chase period.

**Fig. 8.**
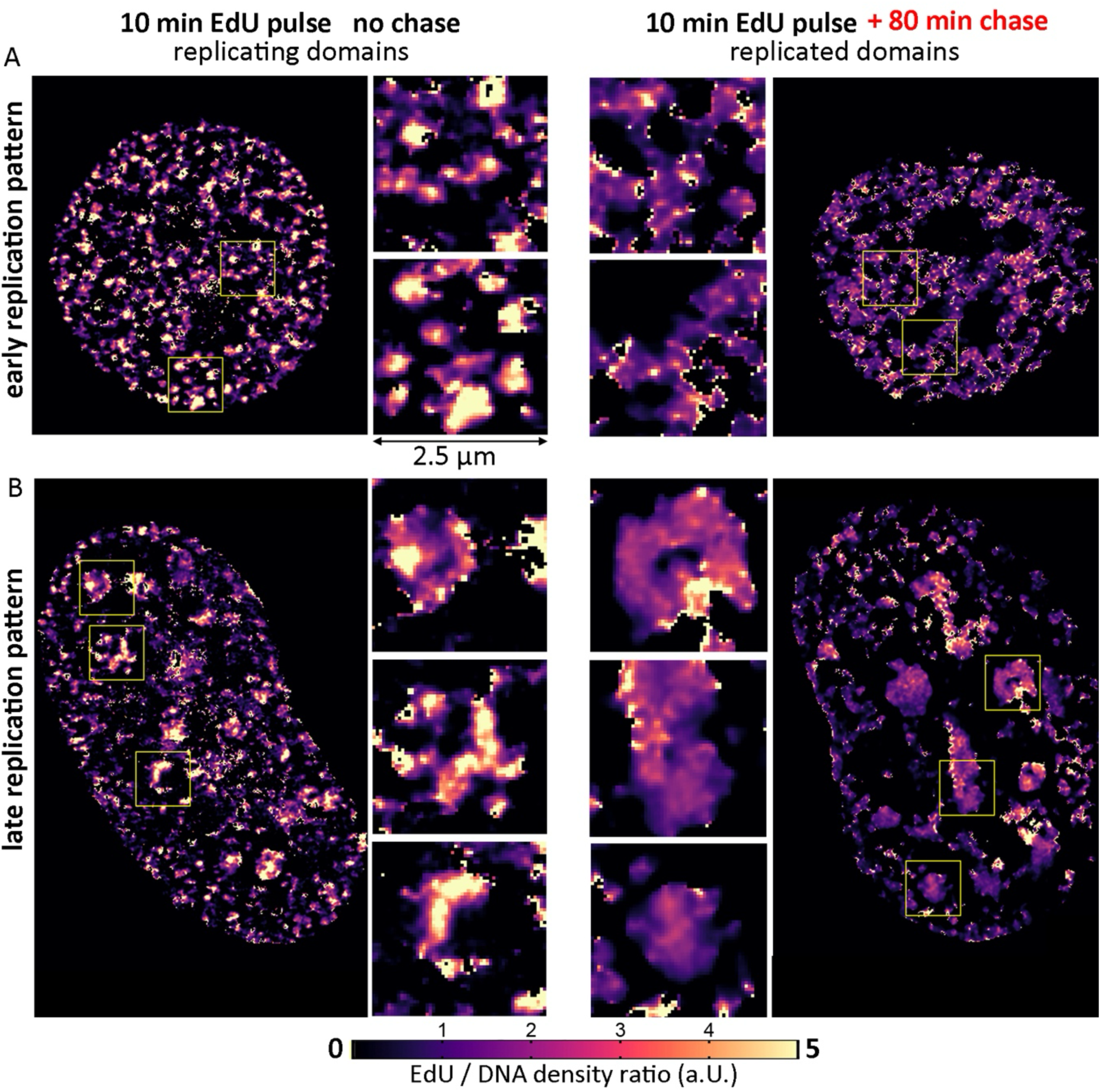
Color-coded ratio images of normalized EdU signals and DNA signal intensities. **(A-B)** Light optical mid-plane sections of the same nuclei as in Fig 4 and 5 are presented as color-coded ratio images of normalized EdU signals and DNA signal intensities. Ratio 1 (dark purple) indicates an amount of EdU signals proportional to the amount of DNA. Ratios 2 to ≥5 indicate the respective excess of EdU labeled DNA. Note clusters with increased ratios >4 (yellow/white) surrounded by EdU-labeled regions with ratios of 1-3 (purple-red) in nuclei of cells fixed immediately aft the 10 min EdU pulse (left). These clusters were much less pronounced in nuclei studied after the 80 min chase (right). Distinct white dots at the periphery of EdU-labeled early and late replicated regions indicate the presence little sites with a fivefold excess of EdU labeled DNA. We interprete these observations as an indication of the not yet completed process of redistribution of EdU-labeled DNA required to restore the original DNA landscape. After completion we would expect a rather uniform pseudo-coloring of EdU-labelled regions close to ratio 1 (deep purple). Acquisitions of the whole nuclei are provided in Supplemental Movies S_movie_5-8

## Discussion

With this study we tested the hypothesis that genome replication, like transcription, also takes place in the ANC (see Introduction, Fig. 3C). Previous studies on transcriptional topography based on quantitative 3D image analysis of nuclei in human, mouse and bovine cell types recorded with 3D SIM had consistently demonstrated a relative over-representation of RNAPII and of epigenetic marks for active chromatin in low DNA density classes, attributed to the ANC, and an under-representation in high DNA intensity classes, attributed to the INC ^36,37,44,46–48^ (for an example, see Fig. 3E). Unexpectedly, replication pulse-labeling experiments (10 min EdU) with replicating CDs demonstrated a relative under-representation of EdU-labeled DNA in low intensity classes 1-2, and an over-representation in high density classes 5-7 (Fig. 7). This disproportional distribution became even more prominent after an additional 80 min chase and emphasizes a dynamic process between these two time points. A decisive difference between transcription and replication of DNA arises from the fact that only a very limited set of DNA sequences is involved in active transcription within the ANC at any given time point. Such sequences are likely to remain there for a considerable time even in case of transcription going on in bursts ^55^. Relocation of coding and regulatory sequences into the compact core region of a CD is considered only for cases, where genes are permanently shut off. In contrast, the requirements of replication where each sequence must be replicated once within a given time window are strongly different.

At face value, our results suggest that DNA replication is accomplished on site also within INC-chromatin. A similar observation was reported in a recent paper of the group of Cristina Cardoso ^56^, where quantitative mapping of EdU labeled replication foci on chromatin compaction classes in different cell types revealed a substantial fraction of early replicating DNA in higher compaction classes. Similarly, Miron et al.^37^ showed a remarkable co-localization of early replicating chromatin with H3M20me3, a histone marker assigned to heterochromatin ^57^. Still, the relative under-representation of EdU signals in low DNA intensity classes does not exclude ongoing replication occurring in the ANC, in line with previously published data that late replicating chromatin in the large heterochromatin blocks of chromocenters recorded by super-resolved images was preferentially noted in rather decondensed channels pervading these blocks ^37,58^ and recently also in plant cells ^59^. Notably, wide IC-channels pervading chromocenters provide particularly fitting experimental situations to study the replication of repetitive DNA within decondensed DNA (ANC) and their re-location into heterochromatin lining these channels (Figs. 5 and 6). Arguably, similar movements take place in chromatin domains with regard to the replication of repressed core regions (Fig. 2). Further studies are necessary to prove or falsify this hypothesis.

In the following, we attempt to integrate our current observations into the context of current knowledge about the 4D nucleome based on a critical assessment of the limited spatial resolution with 3D SIM, and the limited time resolution of the entire genome replication process achieved with an EdU-pulse labeling of 10 min and a chase period of 80 min. Several possibilities can be considered to explain the apparent execution of DNA replication within the INC. First, heterochromatin may be less densely compacted than suggested by ^29^ thus enabling the penetration of replisome complexes formed in the IC-channels into zonal regions with increasingly compact chromatin (Fig. 2C-D). This explanation would be in line with a relatively open configuration of CDs (Fig. 2B). Second, an interpretation could, however, also be reconciled with the CD model predicting a highly compact core region surrounded by decondensed chromatin ^29,37^ (Fig. 2A, C), with additional assumptions on the space-time dynamics during and after CD replication. A replicon may be considered responsible for the replication a chromatin nanodomain (CND) of about 100 kb, (for review see ^60^) located in the core region of such a CD. Local decondensation of such a CND would not solve the accessibility problem of a replisome complex formed outside the CD in case that the CND is still surrounded by a rim of highly compact chromatin, but leaves the option that individual components of such a machine move into the core region and form the replisome complex inside. Alternatively, the whole CND may be moved into the periphery of the CD, forming a replicon there composed of decondensed chromatin. With a lateral resolution of ~120 nm and an axial resolution of ~300 nm, 3D-SIM lacks the spatial resolution to explore chromatin substructures within CDs. The true size and compaction of a CND or even a smaller sized nucleosome clutch (<10 kb) ^27,28^ (for review see^60^) with a physical configuration below the resolution limit cannot be measured. With 3D SIM, we can dissect only the formation of EdU labeled clusters with sizes exceeding the resolution limit resulting from a spatial accumulation of numerous submicroscopic events of chromatin duplication.

In addition to the limited spatial resolution of 3D SIM, the observation of details of genome replication within CDs is also limited by a lack of time resolution of snap-shots of nuclei recorded and quantitatively assessed after two time points, either fixed immediately after a 10 min pulse or after an additional 80 min chase. Both shorter and longer time points need to be studied for a more detailed insight into the dynamic movements of chromatin involved in CD replication. Under the assumption that replication of CDs occurs along decondensed stretches of nucleosome clutches, followed by rapid compaction and further that the replicated nucleosome clutches are closely spaced and form larger compacted aggregates already within a pulse-labelling period of 10 min, they will be detected in high density classes attributed to the INC despite all ongoing replication taking place in decondensed chromatin. This could explain our observation of an overrepresentation of EdU signals in high density DNA classes in “snapshots” of early replicating nuclei recorded with 3D SIM already after 10 min pulse-labelling notwithstanding a necessity for a rapid, transient chromatin decondensation at sites of ongoing replication. A replication fork acts at both sides of a replicon with a speed of 2-3kbp/min of newly replicated DNA ^25^. Accordingly, the replication of a 10 kb nucleosome clutch may require less than 5 min. Even with higher microscopic resolution, a 10 min EdU pulse may be too long to confirm or exclude a rapid local decompaction and recompaction of nucleosome clutches.

A chase time of 80 min was likely sufficient for completion of DNA replication of pulse-labeled CDs, but apparently too short for a complete restoration of the replicated CDs within the context of the nuclear landscape (Fig. 2D, E, Fig. 7). Understanding the spatial dynamics of labeled chromatin awaits further studies with both improved spatial and time resolution, ideally live cell approaches ^61^. Arguably, condensed chromatin is the default chromatin state in higher eukaryotic cells ^31^. In line with this, replication of a mother CD with a condensed core region and transcriptionally active periphery (Fig. 2C) may initially yield two largely inactive daughter CDs. This default state requires an ATP-dependent process for the reformation of transcriptionally active loops, likely including ATP-dependent remodeling complexes like SWI/SNF ^62^. Apparently, the restructuring of each replicated CDs takes much longer than the time of ~1 hour needed for DNA replication. For more detailed insights into the entire dynamic process, CD replication microscopy with higher resolution than 3D SIM is required to study genome replication at the level of nucleosome clutches of individual replicons. It also essential to visualize the replication machinery together with replicating DNA, as well as other machineries involved in the reconfiguration of replicated chromatin. Ideally, one would like to study these events live in individual cells. Recently developed Hi-C methods, such as sister-chromatid-sensitive Hi-C, where Hi-C contacts can be classified as cis- or trans-sister contact, enable the discrimination of sister chromatids in replicated chromosomes ^63^. This opens a way to reveal how sister chromatids fold relative to each other and enable investigations on the conformation of replicated chromosomes.

For a comprehensive understanding of genome replication, it is necessary to understand the space-dynamics of an interphase chromatin organization that allows the apparently instantaneous formation of mitotic chromosomes with two sister chromatids at the onset of prophase required for the faithful separation of chromatids during anaphase^64^ and the rapid reformation of functional 4D nucleomes in daughter cells. The spatial separation of daughter CDs arranged along the two sister chromatids formed in each CT during S-phase likely starts as soon as such a pair is formed. Separation of sister chromatids in G2-CTs precedes the rapid formation of mitotic chromosomes with distinct sister chromatids ^65–67^. According to a recent study, the loop-extruding activity of cohesin is required to resolve sister chromatids during G2 ^68^. Cohesin depletion, however, does not prevent cells to enter mitosis forming mitotic chromosomes with distinct chromatids able to perform chromatid segregation, though frequently with misalignment ^36^.

## Acknowledgments

PMI28 cells were kindly provided by Irina Solovei (Biocenter LMU), the hybrid *Mus musculus* × *Mus castaneus* female ESC line 16.7 was kindly provided by J. Lee, Harvard Medical School. We thank Sandra Ritz and all the members of the IMB Microscopy Core Facility for technical support. We thank Christoph Drees for support with the Elyra 7 microscope. The Zeiss Elyra was funded by the Gutenberg Research College and Deutsche Forschungsgemeinschaft (DFG, German Research Foundation) – 497845157.

## List of abbreviations

ANC: active nuclear compartment
CD: chromatin domain
CND: chromatin nanodomain
CT: chromosome territory
ESC: embryonic stem cells
INC: inactive nuclear compartment
Mb(p): Megabase (pair)
ND: nucleosomal domain
RD: replication domain
RNAPII: RNA polymerase II
TAD: topologically associating domain

## Supplementary movies (on request to M. Cremer)

S_movie 1. Whole acquisition of nucleus shown in Fig. 4A (early replication, 10min EdU pulse)

S_movie 2. Whole acquisition of nucleus shown in Fig. 4B (early replication, 10min EdU pulse+80min chase)

S_movie 3. Whole acquisition of nucleus shown in Fig. 5A (late replication, 10min EdU pulse)

S_movie 4. Whole acquisition of nucleus shown in Fig. 5B (late replication, 10min EdU pulse+80min chase)

S_movie 5. Whole acquisition of nucleus shown in Fig. 8A left (ratio image, early replication, 10min EdU pulse)

S_movie 6. Whole acquisition of nucleus shown in Fig. 8A right (ratio image, early replication, 10min EdU pulse+80min chase)

S_movie 7. Whole acquisition of nucleus shown in Fig. 8B left (ratio image, late replication, 10min EdU pulse)

S_movie 8. Whole acquisition of nucleus shown in Fig. 8B right (ratio image, late replication, 10min EdU pulse+80min chase)

## Notes

### Competing Interest Statement

The authors have declared no competing interest.

## References

1. Cremer, T., and Cremer, C. (2001). Chromosome territories, nuclear architecture and gene regulation in mammalian cells. Nat Rev Genet 2, 292–301. 10.1038/35066075.

2. Cremer, T., Cremer, M., Hubner, B., Strickfaden, H., Smeets, D., Popken, J., Sterr, M., Markaki, Y., Rippe, K., and Cremer, C. (2015). The 4D nucleome: Evidence for a dynamic nuclear landscape based on co-aligned active and inactive nuclear compartments. FEBS Lett 589, 2931–2943. 10.1016/j.febslet.2015.05.037.

3. Dekker, J., Alber, F., Aufmkolk, S., Beliveau, B.J., Bruneau, B.G., Belmont, A.S., Bintu, L., Boettiger, A., Calandrelli, R., Disteche, C.M., et al. (2023). Spatial and temporal organization of the genome: Current state and future aims of the 4D nucleome project. Mol Cell 83, 2624–2640. 10.1016/j.molcel.2023.06.018.

4. Dekker, J., Belmont, A.S., Guttman, M., Leshyk, V.O., Lis, J.T., Lomvardas, S., Mirny, L.A., O’Shea, C.C., Park, P.J., Ren, B., et al. (2017). The 4D nucleome project. Nature 549, 219–226. 10.1038/nature23884.

5. Di Stefano, M., Paulsen, J., Jost, D., and Marti-Renom, M.A. (2021). 4D nucleome modeling. Curr Opin Genet Dev 67, 25–32. 10.1016/j.gde.2020.10.004.

6. Rey-Millet, M., Bystricky, K., and International Nucleome, C. (2024). The genome in space and time comes of age. Nucleus 15, 2307665. 10.1080/19491034.2024.2307665.

7. Soroczynski, J., and Risca, V.I. (2023). Technological advances in probing 4D genome organization. Curr Opin Cell Biol 84, 102211. 10.1016/j.ceb.2023.102211.

8. Tashiro, S., and Lanctot, C. (2015). The International Nucleome Consortium. Nucleus 6, 89–92. 10.1080/19491034.2015.1022703.

9. Attali, I., Botchan, M.R., and Berger, J.M. (2021). Structural Mechanisms for Replicating DNA in Eukaryotes. Annu Rev Biochem 90, 77–106. 10.1146/annurev-biochem-090120-125407.

10. Burgers, P.M.J., and Kunkel, T.A. (2017). Eukaryotic DNA Replication Fork. Annu Rev Biochem 86, 417–438. 10.1146/annurev-biochem-061516-044709.

11. Marchal, C., Sima, J., and Gilbert, D.M. (2019). Control of DNA replication timing in the 3D genome. Nat Rev Mol Cell Biol 20, 721–737. 10.1038/s41580-019-0162-y.

12. Mechali, M. (2010). Eukaryotic DNA replication origins: many choices for appropriate answers. Nat Rev Mol Cell Biol 11, 728–738. 10.1038/nrm2976.

13. Zhao, P.A., Rivera-Mulia, J.C., and Gilbert, D.M. (2017). Replication Domains: Genome Compartmentalization into Functional Replication Units. Adv Exp Med Biol 1042, 229–257. 10.1007/978-981-10-6955-0_11.

14. Zhang, W., Feng, J., and Li, Q. (2020). The replisome guides nucleosome assembly during DNA replication. Cell Biosci 10, 37. 10.1186/s13578-020-00398-z.

15. Jones, M.L., Baris, Y., Taylor, M.R.G., and Yeeles, J.T.P. (2021). Structure of a human replisome shows the organisation and interactions of a DNA replication machine. EMBO J 40, e108819. 10.15252/embj.2021108819.

16. Reinhart, M., and Cardoso, M.C. (2017). A journey through the microscopic ages of DNA replication. Protoplasma 254, 1151–1162. 10.1007/s00709-016-1058-8.

17. Taylor, J.H., and Hozier, J.C. (1976). Evidence for a four micron replication unit in CHO cells. Chromosoma 57, 341–350. 10.1007/BF00332159.

18. Boulos, R.E., Drillon, G., Argoul, F., Arneodo, A., and Audit, B. (2015). Structural organization of human replication timing domains. FEBS Lett 589, 2944–2957. 10.1016/j.febslet.2015.04.015.

19. Dimitrova, D.S., and Berezney, R. (2002). The spatio-temporal organization of DNA replication sites is identical in primary, immortalized and transformed mammalian cells. J Cell Sci 115, 4037–4051. 10.1242/jcs.00087.

20. Alexandrova, O., Solovei, I., Cremer, T., and David, C.N. (2003). Replication labeling patterns and chromosome territories typical of mammalian nuclei are conserved in the early metazoan Hydra. Chromosoma 112, 190–200. 10.1007/s00412-003-0259-z.

21. Postberg, J., Alexandrova, O., Cremer, T., and Lipps, H.J. (2005). Exploiting nuclear duality of ciliates to analyse topological requirements for DNA replication and transcription. J Cell Sci 118, 3973–3983. 10.1242/jcs.02497.

22. Rhind, N., and Gilbert, D.M. (2013). DNA replication timing. Cold Spring Harb Perspect Biol 5, a010132. 10.1101/cshperspect.a010132.

23. Jackson, D.A., and Pombo, A. (1998). Replicon clusters are stable units of chromosome structure: evidence that nuclear organization contributes to the efficient activation and propagation of S phase in human cells. J Cell Biol 140, 1285–1295.

24. Schermelleh, L., Solovei, I., Zink, D., and Cremer, T. (2001). Two-color fluorescence labeling of early and mid-to-late replicating chromatin in living cells. Chromosome Res 9, 77–80.

25. Chagin, V.O., Casas-Delucchi, C.S., Reinhart, M., Schermelleh, L., Markaki, Y., Maiser, A., Bolius, J.J., Bensimon, A., Fillies, M., Domaing, P., et al. (2016). 4D Visualization of replication foci in mammalian cells corresponding to individual replicons. Nat Commun 7, 11231. 10.1038/ncomms11231.

26. Xiang, W., Roberti, M.J., Heriche, J.K., Huet, S., Alexander, S., and Ellenberg, J. (2018). Correlative live and super-resolution imaging reveals the dynamic structure of replication domains. J Cell Biol 217, 1973–1984. 10.1083/jcb.201709074.

27. Otterstrom, J., Castells-Garcia, A., Vicario, C., Gomez-Garcia, P.A., Cosma, M.P., and Lakadamyali, M. (2019). Super-resolution microscopy reveals how histone tail acetylation affects DNA compaction within nucleosomes in vivo. Nucleic Acids Res 47, 8470–8484. 10.1093/nar/gkz593.

28. Ricci, M.A., Manzo, C., Garcia-Parajo, M.F., Lakadamyali, M., and Cosma, M.P. (2015). Chromatin fibers are formed by heterogeneous groups of nucleosomes in vivo. Cell 160, 1145–1158. 10.1016/j.cell.2015.01.054.

29. Gelleri, M., Chen, S.Y., Hubner, B., Neumann, J., Kroger, O., Sadlo, F., Imhoff, J., Hendzel, M.J., Cremer, M., Cremer, T., et al. (2023). True-to-scale DNA-density maps correlate with major accessibility differences between active and inactive chromatin. Cell Rep 42, 112567. 10.1016/j.celrep.2023.112567.

30. Mansisidor, A.R., and Risca, V.I. (2022). Chromatin accessibility: methods, mechanisms, and biological insights. Nucleus 13, 236–276. 10.1080/19491034.2022.2143106.

31. Maeshima, K., Iida, S., Shimazoe, M.A., Tamura, S., and Ide, S. (2024). Is euchromatin really open in the cell? Trends Cell Biol 34, 7–17. 10.1016/j.tcb.2023.05.007.

32. Maeshima, K., Kaizu, K., Tamura, S., Nozaki, T., Kokubo, T., and Takahashi, K. (2015). The physical size of transcription factors is key to transcriptional regulation in chromatin domains. J Phys Condens Matter 27, 064116. 10.1088/0953-8984/27/6/064116.

33. Demmerle, J., Hao, S., and Cai, D. (2023). Transcriptional condensates and phase separation: condensing information across scales and mechanisms. Nucleus 14, 2213551. 10.1080/19491034.2023.2213551.

34. Cremer, T., Cremer, M., Hubner, B., Silahtaroglu, A., Hendzel, M., Lanctot, C., Strickfaden, H., and Cremer, C. (2020). The Interchromatin Compartment Participates in the Structural and Functional Organization of the Cell Nucleus. Bioessays 42, e1900132. 10.1002/bies.201900132.

35. Nozaki, T., Shinkai, S., Ide, S., Higashi, K., Tamura, S., Shimazoe, M.A., Nakagawa, M., Suzuki, Y., Okada, Y., Sasai, M., et al. (2023). Condensed but liquid-like domain organization of active chromatin regions in living human cells. Sci Adv 9, eadf1488. 10.1126/sciadv.adf1488.

36. Cremer, M., Brandstetter, K., Maiser, A., Rao, S.S.P., Schmid, V.J., Guirao-Ortiz, M., Mitra, N., Mamberti, S., Klein, K.N., Gilbert, D.M., et al. (2020). Cohesin depleted cells rebuild functional nuclear compartments after endomitosis. Nat Commun 11, 6146. 10.1038/s41467-020-19876-6.

37. Miron, E., Oldenkamp, R., Brown, J.M., Pinto, D.M.S., Xu, C.S., Faria, A.R., Shaban, H.A., Rhodes, J.D.P., Innocent, C., de Ornellas, S., et al. (2020). Chromatin arranges in chains of mesoscale domains with nanoscale functional topography independent of cohesin. Science Advances 6, eaba8811. DOI: 10.1126/sciadv.aba8811

38. Strickfaden, H., Tolsma, T.O., Sharma, A., Underhill, D.A., Hansen, J.C., and Hendzel, M.J. (2020). Condensed Chromatin Behaves like a Solid on the Mesoscale In Vitro and in Living Cells. Cell 183, 1772–1784 e1713. 10.1016/j.cell.2020.11.027.

39. Hubner, B., Strickfaden, H., Muller, S., Cremer, M., and Cremer, T. (2009). Chromosome shattering: a mitotic catastrophe due to chromosome condensation failure. Eur Biophys J 38, 729–747. 10.1007/s00249-009-0496-z.

40. Hancock, R. (2004). A role for macromolecular crowding effects in the assembly and function of compartments in the nucleus. J Struct Biol 146, 281–290. 10.1016/j.jsb.2003.12.008.

41. Hancock, R. (2014). The crowded nucleus. Int Rev Cell Mol Biol 307, 15–26. 10.1016/B978-0-12-800046-5.00002-3.

42. Hancock, R. (2018). Crowding, Entropic Forces, and Confinement: Crucial Factors for Structures and Functions in the Cell Nucleus. Biochemistry (Mosc) 83, 326–337. 10.1134/S0006297918040041.

43. Iida, S., Ide, S., Tamura, S., Tani, T., Goto, T., Shribak, M., and Maeshima, K. (2024). Orientation-Independent-DIC imaging reveals that a transient rise in depletion force contributes to mitotic chromosome condensation. bioRxiv preprint. 10.1101/2023.11.11.566679

44. Smeets, D., Markaki, Y., Schmid, V.J., Kraus, F., Tattermusch, A., Cerase, A., Sterr, M., Fiedler, S., Demmerle, J., Popken, J., et al. (2014). Three-dimensional super-resolution microscopy of the inactive X chromosome territory reveals a collapse of its active nuclear compartment harboring distinct Xist RNA foci. Epigenetics Chromatin 7, 8. 10.1186/1756-8935-7-8.

45. Fakan, S., and van Driel, R. (2007). The perichromatin region: a functional compartment in the nucleus that determines large-scale chromatin folding. Semin Cell Dev Biol 18, 676–681. 10.1016/j.semcdb.2007.08.010.

46. Cremer, M., Schmid, V.J., Kraus, F., Markaki, Y., Hellmann, I., Maiser, A., Leonhardt, H., John, S., Stamatoyannopoulos, J., and Cremer, T. (2017). Initial high-resolution microscopic mapping of active and inactive regulatory sequences proves non-random 3D arrangements in chromatin domain clusters. Epigenetics Chromatin 10, 39. 10.1186/s13072-017-0146-0.

47. Hubner, B., Lomiento, M., Mammoli, F., Illner, D., Markaki, Y., Ferrari, S., Cremer, M., and Cremer, T. (2015). Remodeling of nuclear landscapes during human myelopoietic cell differentiation maintains co-aligned active and inactive nuclear compartments. Epigenetics Chromatin 8, 47. 10.1186/s13072-015-0038-0.

48. Popken, J., Brero, A., Koehler, D., Schmid, V.J., Strauss, A., Wuensch, A., Guengoer, T., Graf, A., Krebs, S., Blum, H., et al. (2014). Reprogramming of fibroblast nuclei in cloned bovine embryos involves major structural remodeling with both striking similarities and differences to nuclear phenotypes of in vitro fertilized embryos. Nucleus 5, 555–589. 10.4161/19491034.2014.979712.

49. Chan, B., and Rubinstein, M. (2023). Activity-driven chromatin organization during interphase: compaction, segregation, and entanglement suppression. bioRxiv preprint. 10.1101/2024.01.22.576729.

50. Schmid, V.J., Cremer, M., and Cremer, T. (2017). Quantitative analyses of the 3D nuclear landscape recorded with super-resolved fluorescence microscopy. Methods 123, 33–46. 10.1016/j.ymeth.2017.03.013.

51. Pau, G., Fuchs, F., Sklyar, O., Boutros, M., and Huber, W. (2010). EBImage--an R package for image processing with applications to cellular phenotypes. Bioinformatics 26, 979–981. 10.1093/bioinformatics/btq046 btq046 [pii].

52. 52. da Costa-Nunes, J.A., Gierlinski, M., Sasaki, T., Haagensen, E.J., Gilbert, D.M., and Blow, J.J. (2023). The location and development of Replicon Cluster Domains in early replicating DNA. Wellcome Open Res 8, 158. 10.12688/wellcomeopenres.18742.2.

53. Heinz, K.S., Rapp, A., Casas-Delucchi, C.S., Lehmkuhl, A., Romero-Fernandez, I., Sanchez, A., Kramer, O.H., Marchal, J.A., and Cardoso, M.C. (2019). DNA replication dynamics of vole genome and its epigenetic regulation. Epigenetics Chromatin 12, 18. 10.1186/s13072-019-0262-0.

54. Wu, R., Singh, P.B., and Gilbert, D.M. (2006). Uncoupling global and fine-tuning replication timing determinants for mouse pericentric heterochromatin. J Cell Biol 174, 185–194. 10.1083/jcb.200601113.

55. Dar, R.D., Razooky, B.S., Singh, A., Trimeloni, T.V., McCollum, J.M., Cox, C.D., Simpson, M.L., and Weinberger, L.S. (2012). Transcriptional burst frequency and burst size are equally modulated across the human genome. Proc Natl Acad Sci U S A 109, 17454–17459. 10.1073/pnas.1213530109.

56. Pradhan, S., and Cardoso, M.C. (2024). Developmental Changes in Genome Replication Progression in Pluripotent versus Differentiated Human Cells. Genes. 10.3390/genes15030305.

57. Ren, W., Fan, H., Grimm, S.A., Kim, J.J., Li, L., Guo, Y., Petell, C.J., Tan, X.F., Zhang, Z.M., Coan, J.P., et al. (2021). DNMT1 reads heterochromatic H4K20me3 to reinforce LINE-1 DNA methylation. Nat Commun 12, 2490. 10.1038/s41467-021-22665-4.

58. Chagin, V.O., Reinhart, B., Becker, A., Mortusewicz, O., Jost, K.L., Rapp, A., Leonhardt, H., and Cardoso, M.C. (2019). Processive DNA synthesis is associated with localized decompaction of constitutive heterochromatin at the sites of DNA replication and repair. Nucleus 10, 231–253. 10.1080/19491034.2019.1688932.

59. Arifulin, E.A., Sorokin, D.V., Anoshina, N.A., Kuznetsova, M., Vayaeva, A.A., Fedotova, A.V., Schubert, V., Kolesnikova, T.D., and Sheval, E.V. (2023). Global nuclear reorganization during heterochromatin replication in the giant-genome plant Nigella damascena L. bioRxiv preprint. 10.1101/2023.08.15.552960.

60. Di Stefano, M., and Cavalli, G. (2022). Integrative studies of 3D genome organization and chromatin structure. Curr Opin Struct Biol 77, 102493. 10.1016/j.sbi.2022.102493.

61. Laine, R.F., Heil, H.S., Coelho, S., Nixon-Abell, J., Jimenez, A., Wiesner, T., Martinez, D., Galgani, T., Regnier, L., Stubb, A., et al. (2023). High-fidelity 3D live-cell nanoscopy through data-driven enhanced super-resolution radial fluctuation. Nat Methods 20, 1949–1956. 10.1038/s41592-023-02057-w.

62. Centore, R.C., Sandoval, G.J., Soares, L.M.M., Kadoch, C., and Chan, H.M. (2020). Mammalian SWI/SNF Chromatin Remodeling Complexes: Emerging Mechanisms and Therapeutic Strategies. Trends Genet 36, 936–950. 10.1016/j.tig.2020.07.011.

63. Mitter, M., and Gerlich, D.W. (2021). Mapping Sister Chromatid Conformation in Replicated Chromosomes. Trends Biochem Sci 46, 169–170. 10.1016/j.tibs.2020.11.011.

64. Sedat, J., McDonald, A., Cang, H., Lucas, J., Arigovindan, M., Kam, Z., Murre, C., and Elbaum, M. (2022). A proposed unified interphase nucleus chromosome structure: Preliminary preponderance of evidence. Proc Natl Acad Sci U S A 119, e2119101119. 10.1073/pnas.2119101119.

65. Eykelenboom, J.K., Gierlinski, M., Yue, Z., Hegarat, N., Pollard, H., Fukagawa, T., Hochegger, H., and Tanaka, T.U. (2019). Live imaging of marked chromosome regions reveals their dynamic resolution and compaction in mitosis. J Cell Biol 218, 1531–1552. 10.1083/jcb.201807125.

66. Ono, T., Yamashita, D., and Hirano, T. (2013). Condensin II initiates sister chromatid resolution during S phase. J Cell Biol 200, 429–441. 10.1083/jcb.201208008.

67. Stanyte, R., Nuebler, J., Blaukopf, C., Hoefler, R., Stocsits, R., Peters, J.M., and Gerlich, D.W. (2018). Dynamics of sister chromatid resolution during cell cycle progression. J Cell Biol 217, 1985–2004. 10.1083/jcb.201801157.

68. Batty, P., Langer, C.C., Takacs, Z., Tang, W., Blaukopf, C., Peters, J.M., and Gerlich, D.W. (2023). Cohesin-mediated DNA loop extrusion resolves sister chromatids in G2 phase. EMBO J 42, e113475. 10.15252/embj.2023113475.

69. Zhang, Y., Boninsegna, L., Yang, M., Misteli, T., Alber, F., and Ma, J. (2024). Computational methods for analysing multiscale 3D genome organization. Nat Rev Genet 25, 123–141. 10.1038/s41576-023-00638-1.

